# Improved models for the relationship between age and the probability of trypanosome infection in female tsetse, *Glossina pallidipes* Austen

**DOI:** 10.1101/2022.09.17.508379

**Authors:** J. W. Hargrove, J. Van Sickle

## Abstract

Between 1990 and 1999, at Rekomitjie Research Station, Zambezi Valley, Zimbabwe, 29,360 female *G. pallidipes* were dissected to determine their ovarian category and trypanosome infection status. Overall prevalences were 3.45% and 2.66% for *T. vivax* and *T. congolense*, respectively, declining during each year as temperatures increased from July - December. Fits to age-prevalence data using Susceptible-Exposed-Infective (SEI) and SI compartmental models were statistically better than those obtained using a published catalytic model, which made the unrealistic assumption that no female tsetse survived more than seven ovulations. The improved models require knowledge of fly mortality, estimated separately from ovarian age distributions. Infection rates were not significantly higher for *T. vivax* than for *T. congolense*. For *T. congolense* in field-sampled female *G. pallidipes*, we found no statistical support for a model where the force of infection was higher at the first feed than subsequently. The long survival of adult female tsetse, combined with feeding at intervals ≤ 3 days, ensures that post-teneral feeds, rather than the first feed, play the dominant role in the epidemiology of *T. congolense* infections in *G. pallidipes*. This is supported by estimates that only about 3% of wild hosts at Rekomitjie were harbouring sufficient *T. congolense* to ensure that tsetse feeding off them take an infected meal, so that the probability of ingesting an infected meal is low at every meal.

## Introduction

Animal African Trypanosomiasis continues to be a serious veterinary problem, costing 37 African countries USD 4.5 billion annually (FAO, 2017). The trypanosome species of veterinary interest in eastern and southern Africa are *Trypanosoma vivax* and *T. congolense*. Woolhouse *et al*. (1993) used a simple catalytic function to model age-related changes in the prevalence of *T. vivax* infections in female *G. pallidipes*. For *T. congolense*, they suggested a slightly more complex model that allowed for a reduction in prevalence among older flies. Age was estimated using ovarian dissection (Challier, 1965), which provides precise estimates of chronological age for flies in their first ovarian cycle – *i.e*., that have ovulated fewer than four times – but not for older flies. Woolhouse *et al*. (1993) assumed that, for their sample, negligible numbers of flies survived to a third ovarian cycle so that uncorrected ovarian category is a valid index of age. This assumption was carried over into other modelling of the same data (Woolhouse & Hargrove, 1998; Lord *et al*., 1999). The larger numbers dissected in the current study makes clear that the above assumption cannot be sustained. Accordingly, we develop improved models that account for the longer survival of adult flies. In so doing, we also investigated the suggestion of Welburn & Maudlin (1992) that the probability of tsetse acquiring *T. congolense* infections was so much lower for bloodmeals other than the first, that non-teneral flies do not play a significant part in the epidemiology of this species of trypanosome. This so called “teneral effect” was based on laboratory work with male and female *G. m. morsitans*; it has not been demonstrated for *G. m. morsitans* in the field, nor for *G. pallidipes* in any setting.

## Methods

### Study area

The study was carried out between September 1990 and April 1999 at Rekomitjie Research Station (16°10’S, 29° 25 E, altitude 520 m), Zambezi Valley, Zimbabwe. Daily maximum and minimum temperatures were recorded using mercury thermometers housed in a Stevenson screen at the Station, and rainfall was recorded using a gauge placed 4 m from the Stevenson screen.

### Fly sampling

Female *G. pallidipes* were collected at various sites within 2 km of the Research Station; >99% of flies were captured in epsilon traps (Hargrove & Langley, 1990) baited with artificial host-odour consisting of acetone, 1-octen-3-ol, 3-n-propyl phenol and 4-methyl phenol released at ~ 200, 0.4, 0.01 and 0.8 mg/h, respectively. The cages in which flies were captured were wrapped in a heavy-duty black cotton cloth and kept in a polystyrene box; in about 90% of cases, traps were cleared at 30-minute intervals. Flies were dissected within 24 hours of capture.

### Classification of flies by ovarian and wing fray category and trypanosome infection status

We followed the general dissection procedure described by Woolhouse et al. (1993). Flies were subjected to ovarian dissection, using the method of Challier (1965), and designated to one of eight ovarian categories (0 - 7) given the relative sizes of ovarioles in the paired ovaries (Hargrove, 2012). Flies in ovarian category 4 include flies that have ovulated 4 + 4*n* times *n* = 0, 1, 2 …. and analogous statements apply to flies in ovarian categories 5, 6 and 7. Flies in ovarian categories 4 – 7 were accordingly assigned ovarian ages of 4 + 4*n* - 7 + 4*n*, respectively.

We used the classification of Jackson (1946) to assign each fly to one of six wing fray categories. While ovarian dissection provides a more accurate estimate of a fly’s chronological age for tsetse, this is only true for flies in ovarian category <4. Wing fray, by contrast, provides a relative measure of age for all flies and allows us to gauge how trypanosome prevalences change over flies’ entire lives.

The mouthparts of each fly were dissected, and the labrum and hypopharynx were examined under a light microscope at 240× magnification. If no trypanosomes were found in the mouthparts the fly was diagnosed as negative for all trypanosome infections and no further dissection was carried out. If trypanosomes were detected, the midgut was dissected out and screened for infection. Where trypanosomes were detected in both the midgut and mouthparts, the fly was diagnosed as being infected with *T. congolense*; if trypanosomes were found only in the mouthparts the fly was diagnosed as being infected with *T. vivax*.

The delay between the time when a fly becomes infected and when it is possible to detect trypanosomes in the fly by dissection is such that there is effectively zero probability of finding trypanosomes in a fly before it has ovulated for the first time. This has been confirmed through the dissection of large numbers of flies in ovarian category 0. Accordingly, flies found to be in ovarian category 0 were not dissected further to look for trypanosomes, and the trypanosome prevalence of such flies was assumed to be identically zero.

### Modelling patterns of acquisition of trypanosome infections

#### Simple catalytic model: assuming no flies ovulate more than seven times

Woolhouse *et al*. (1993) used a simple catalytic model to explain age-related changes in the prevalence of *T*. *vivax* infections in female *G. pallidipes*, fitting their data using the function:

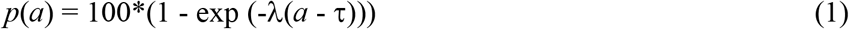

where *p*(*a*) is the trypanosome prevalence among flies of age *a* days, λ the force of infection and τ the number of days between the acquisition of an infection and the time when trypanosomes can first be detected in the fly. Equation (1) predicts prevalence directly for flies of any known daily age, without requiring knowledge of fly mortality, and without using explicit predicted values for the numbers of infected and uninfected flies. We follow Woolhouse *et al*. (1993) in applying analyses to data pooled over time, so that the parameter estimates from the model are viewed as long-term averages.

We applied the catalytic model to our data. Since our data are classified only according to the ovarian category of the flies dissected, it is necessary to assign the daily ages of each ovarian age. Following Hargrove (1995, 2012) we allocate flies of age 0-7 days to category zero: thereafter we assume a 9-day pregnancy cycle so that the endpoints of ovarian ages 1 to 7 occur at ages 16, 25, 34, 43, 52, 61 and 70 days, respectively. The predicted prevalence, *P(i)*, for flies of ovarian age *i*, *i* = 1,7 is then given by:

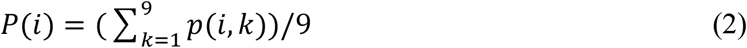

where *p*(*i*, *k*) is the prevalence of a fly of ovarian age *i*, on day *k* of pregnancy. As explained above, we assume *P*(0) = 0.

#### Susceptible-Infected SI model: assuming flies can survive to ovulate more than seven times

If significant proportions of female tsetse survive to ovulate more than seven times, Equations (1) and (2) are inaccurate. This is because it is not possible to differentiate flies that have ovulated 4, 8 or 12 times: nor those that have ovulated 5, 9 or 13 times, 6, 10 or 14 times, or 7, 11 or 15 times (Challier, 1965). These flies are pooled into ovarian categories 4+4*n*, 5+4*n*, 6+4*n* and 7+4*n*, respectively. Field results suggest that few flies will survive to ovulate more than 15 times (Hargrove *et al*., 2011) and we assume this approximation holds for all further calculations in this study.

To handle this complication, we introduce an SI model and track separately the numbers of infected (*I*), noninfected (susceptible, *S*), and total (*N* = *S*+*I*) flies. We also model fly mortality, by calculating the total numbers *N*(*a*) = *N*_0_ exp(-μ*a*) of flies surviving to age *a* days, where μ is the instantaneous mortality rate, with units days^−1^, and related to the daily survival probability by φ = exp(-μ). The parameter *N_0_* is the number in category 0. With this mortality model, the numbers of surviving flies that are infected is given by *I*(*a*) = *N*_0_ exp(-μ*a*)(1 - exp (-λ(*a* - τ)), where λ and τ are as defined for Equation (1). The total numbers, and numbers infected, at each daily age are then pooled, as described above, according to their ovarian age, *i*, *i* = 1, 15. For ovarian categories 1-3, and daily age indexed by *k*, we have:

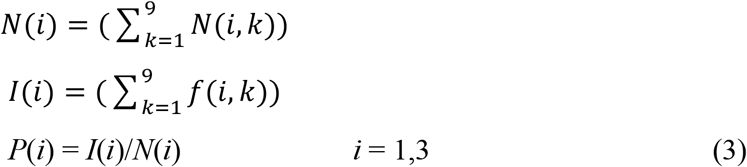

For older flies, we need to carry out the following pooling:

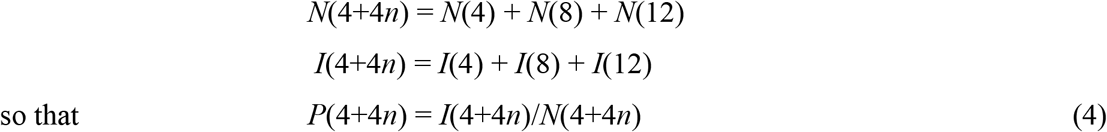

and analogous results apply for flies in ovarian categories 5+4*n*, 6+4*n* and 7+4*n*. Note that the parameter *N_0_* can be an arbitrary value because it divides out when calculating *P(i*).

The SI model requires a knowledge of the daily survival probability φ. We follow Woolhouse *et al*. (1993) in assuming that φ is independent both of a fly’s age and of its infection status and, accordingly, we estimate mortality directly from the ovarian age distribution of all tsetse (see below) and use this same mortality estimate in the SI and SEI models.

#### Susceptible, Exposed and Infective (SEI) model

As an alternative to the SI model, we consider a model where, instead of having a fixed delay term (τ), we partition flies into three states – Susceptible, Exposed and Infective (SEI). Susceptible flies are those that have never acquired a trypanosome infection. Exposed flies have been so infected, but the trypanosome infection has not developed to the point where the fly can pass on that infection – *i.e*., it is not infective. Infective flies can pass on the trypanosome infection, and we assume that flies only leave this state by dying. They are also the only flies in which trypanosome can be detected using dissection techniques; accordingly, only the infective flies are included in compartment I in the modelling. The SI model, with a fixed time delay (τ), is biologically unrealistic in assuming a sharp boundary between the ages when tsetse are and are not infective; this problem does not arise for the SEI model. We assume an arbitrary initial number *S_0_* of susceptible flies. Since none have yet been exposed or infected, *E_0_* = 0 and *I_0_* = 0. Then the following transition equations apply for the SEI model as the flies age:

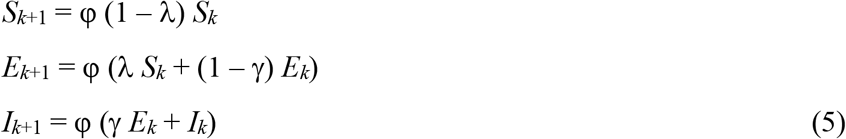

where, for each unit of time, φ now defines the proportion surviving, λ the probability that susceptibles (*S_k_*) move to the exposed class (*E*_*k*+1_), and γ the probability of moving from the exposed class (*E_k_*) to the infective class (*I*_*k*+1_), between time periods *i* and *i*+1. The total count is given by *N*_*k*+1_ = *S*_*k*+1_ + *E*_*k*+1_ + *I*_*k*+1_. When implementing Equations 5 we used the same, separately estimated, value of φ as used in the SI model.

When applying this model, we changed the timestep from 1 to 3 days, assuming that the latter timestep approximated the interval between successive feeds, and that the flies take an average of three meals in a 9-day pregnancy (Randolph *et al*., 1992). Using the feeding interval as the time step is more realistic because flies can only become infected when they are actually taking a bloodmeal: the change also enabled us to investigate the suggestion that the force of infection is higher at the first blood meal than at subsequent meals (Welburn *et al*., 1992).

With a 3-day timestep, there are 3 age increments per 9-day ovarian cycle. Thus, predicted prevalences for the SEI model are calculated with modified versions of Equations 3 and 4. For example, starting at age 7 days, ovarian category 1 includes the 3rd, 4th and 5th age increments, so that

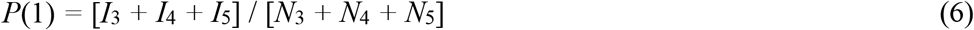

where the subscript indexes 3-day periods, with analogous equations for *P*(2) and *P*(3). For the pooled ovarian categories, we have

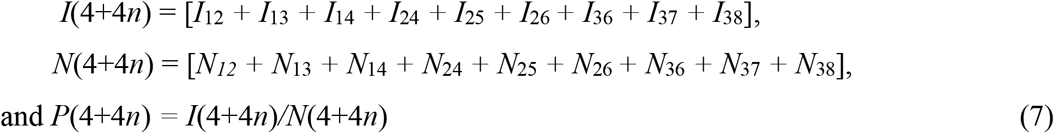

with analogous expressions for *P*(5+4*n*), *P*(6+4*n*) and *P*(7+4*n*).

### Statistical methods

#### Prevalence models: parameter estimation and model assessment

We estimated the parameters of all models using maximum likelihood (ML), as implemented in the R package *maxlik* (Henningsen & Toomet, 2011). For the catalytic, SI and SEI models, the predicted prevalence can also be interpreted as the predicted probability that a randomly sampled fly is infected. Thus, the log likelihood for these three models has the binomial form:

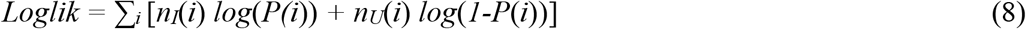

where *n_I_*(*i*) and *n_U_*(*i*)= *n*(*i*) - *n_I_*(*i*) are the observed counts of infected and uninfected flies, respectively, in the 7 ovarian categories *i* = 1, 2, 3, 4+4*n*, .., 7+4*n*. The observed total count in each category is *n*(*i*).

The three prevalence models differ only in how they predict the *P*(*i*) values of Equation 8. For the catalytic model, *P*(*i*) is given by Equation 2, as a function of model parameters λ and τ. Likewise, the predicted *P*(*i*) values are given by Equations 3 and 4 for the SI model, and by Equations 6 and 7 for the SEI model.

We report 95% confidence intervals for all estimated parameters. In addition, we used the Akaike Information Criterion (AIC), as well as graphical displays, to assess and compare how well the prevalence models fit the data (Burnham & Anderson, 2002). AIC measures a model’s quality of fit, with a penalty for the number of estimated parameters needed to achieve that quality. In calculating AIC for the SI and SEI models, we also included φ in the count of estimated parameters. As a rule of thumb, models having AIC values within 2 units of each other do not differ in their quality of fit (Burnham & Anderson, 2002).

#### Estimation of survival probability

We also used ML to estimate the survival probability φ based only on the total counts, and separately from the prevalence models. This estimate was then used as a fixed parameter value when estimating the other parameters of the SI and SEI models. To estimate φ, we assumed that the total counts, *N*(*i*), were a sample from a stationary age distribution of live flies. Then, the number surviving to ovarian age *k* is given by *N*(*i*) = *N_0_* φ*^i^*, where *N_0_* is an arbitrary initial count of age 0 flies. Assuming that no flies survive beyond 15 ovulations, the grand total count of all flies is *N_T_* = *N_0_∑_i_* φ*^i^*, where the sum ranges over *k* = 1,2 … 15. Then, standardization of the *N*(*i*) by *N_T_* expresses the age distribution as a multinomial distribution of the probabilities *p*(*i*) that a randomly-selected fly is of ovarian age *i*.

For ovarian ages 1, 2, 3, the probabilities are:

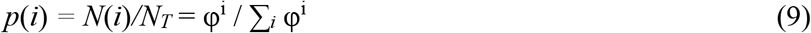

The number of survivors in the pooled ovarian category (4+4*n*) includes all flies that survived through ovarian cycles 4, 8 or 12, and similarly for the other pooled categories. Thus the probabilities for pooled categories are:

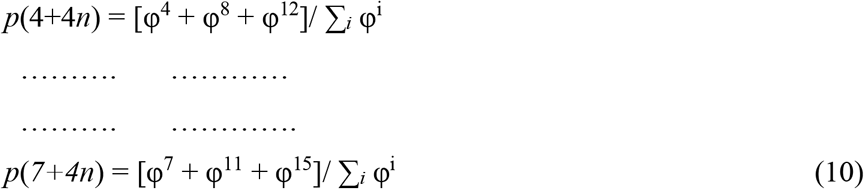

Finally, the log likelihood is given by

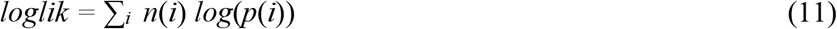

where *n*(*i*) are the observed total counts in ovarian categories 1, 2, .. 4+4*n*, .. 7+4*n*, and the probabilities *p*(*i*) are given by Equations 9 and 10. We maximized Equation 11 directly. Equivalently, Hargove (1993) set the derivative of Equation 11 to zero, thus yielding a closed form for the maximum, which was then solved numerically.

As defined above, φ is the probability of survival for 9 days. We converted the estimate of φ into 3-day and 1-day survival probabilities, for use in the SEI and SI models, by calculating its cube root and ninth root, respectively.

## Results

### Temporal variations in temperature and rainfall, and in trypanosome prevalence

Rekomitjie Research Station has a hot tropical climate with temperatures increasing between July and November, leading up to the start of the single rainy season, largely restricted to November-February (Figure 1A). A total of 29,360 female *G. pallidipes* were dissected for determination of ovarian category and trypanosome infection status. Of these, the wing fray category was gauged in 27,951 flies. *T. vivax* and *T. congolense* infection prevalences – for all flies pooled on age – were 3.45% and 2.66%, respectively and declined during each year as temperatures increased between about July and December. Thereafter they increased sharply for 1-2 months, then more slowly to peak again in July (Figure 1B).

**Figure 1.**
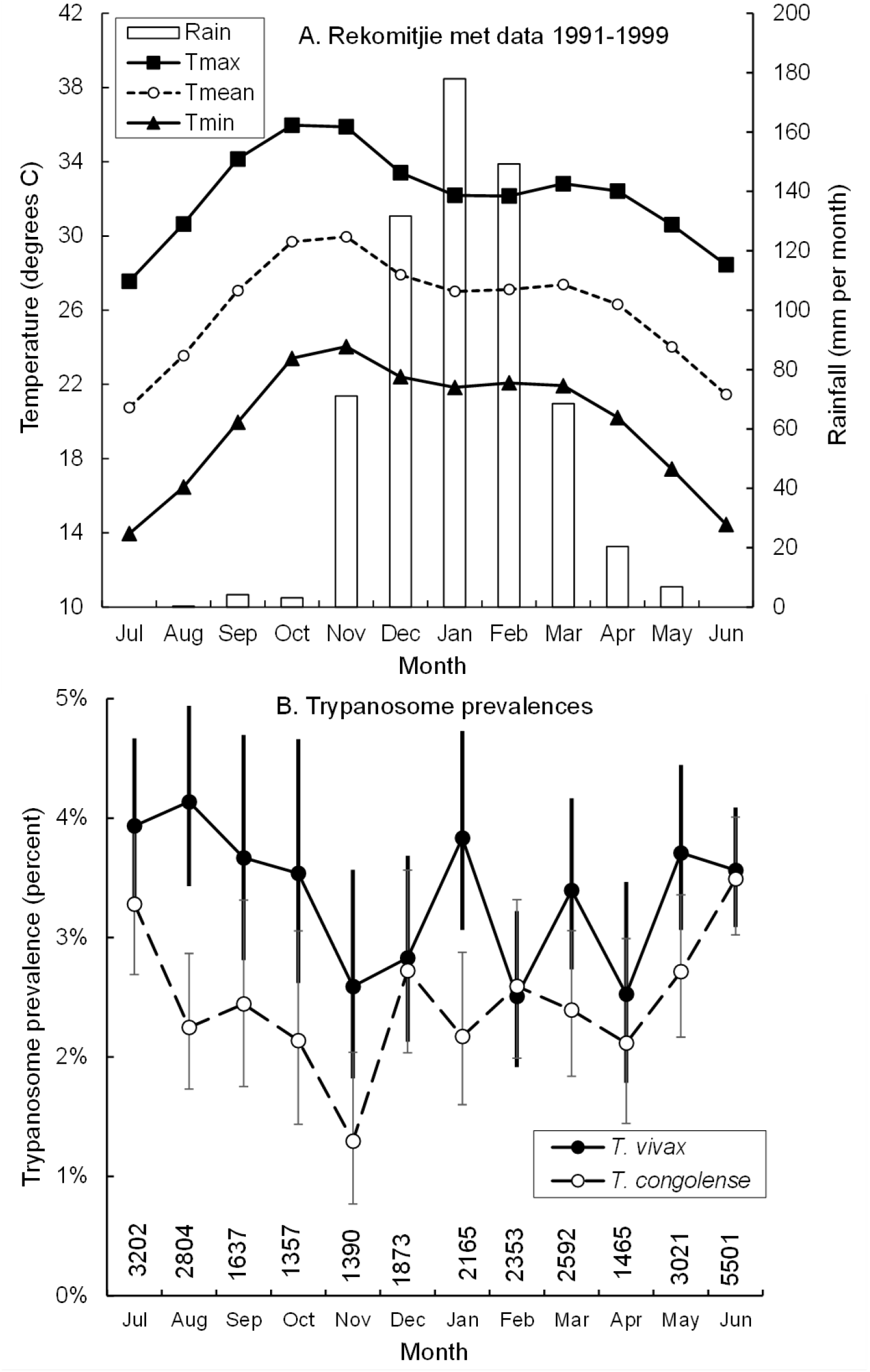
A. Mean daily temperatures and mean monthly rainfall 1991-1999, Rekomitjie Research Station, Zambezi Valley, Zimbabwe. B. Monthly prevalence of infections of *T. vivax* and *T. congolense* in female *G. pallidipes* captured and dissected between 1990 and 1999. Error bars indicate the 95% confidence intervals. Samples sizes are for each month, pooled on year.

### Age distribution of all flies dissected, and of those found infected with trypanosomes

Among all flies dissected, there were similar numbers of ovarian age 1 and 2, smaller numbers in ovarian age 3, and then a marked jump in numbers when moving to ovarian age 4 + 4*n* (Figure 2A) – in accord with the fact that ovarian category 4 includes flies that have ovulated 4, 8, 12 etc times (see above). For infected flies, numbers increase between ages 1 and 3 and then take an even larger jump for those in age group 4 – since flies can accrue infections throughout life (Figure 2B). The numbers of infected flies only begin to diminish for flies of ovarian age 4, when older flies begin to die at enhanced rates (Hargrove et al., 2011). When wing fray was used as a cruder measure of age, the numbers dissected increased until wing fray category 4, and the numbers of infected flies until category 4 or 5 (Figure 2C, D). The difference reflects the fact that wing fray cannot decrease with age – and will naturally increase at different rates in different flies.

**Figure 2.**
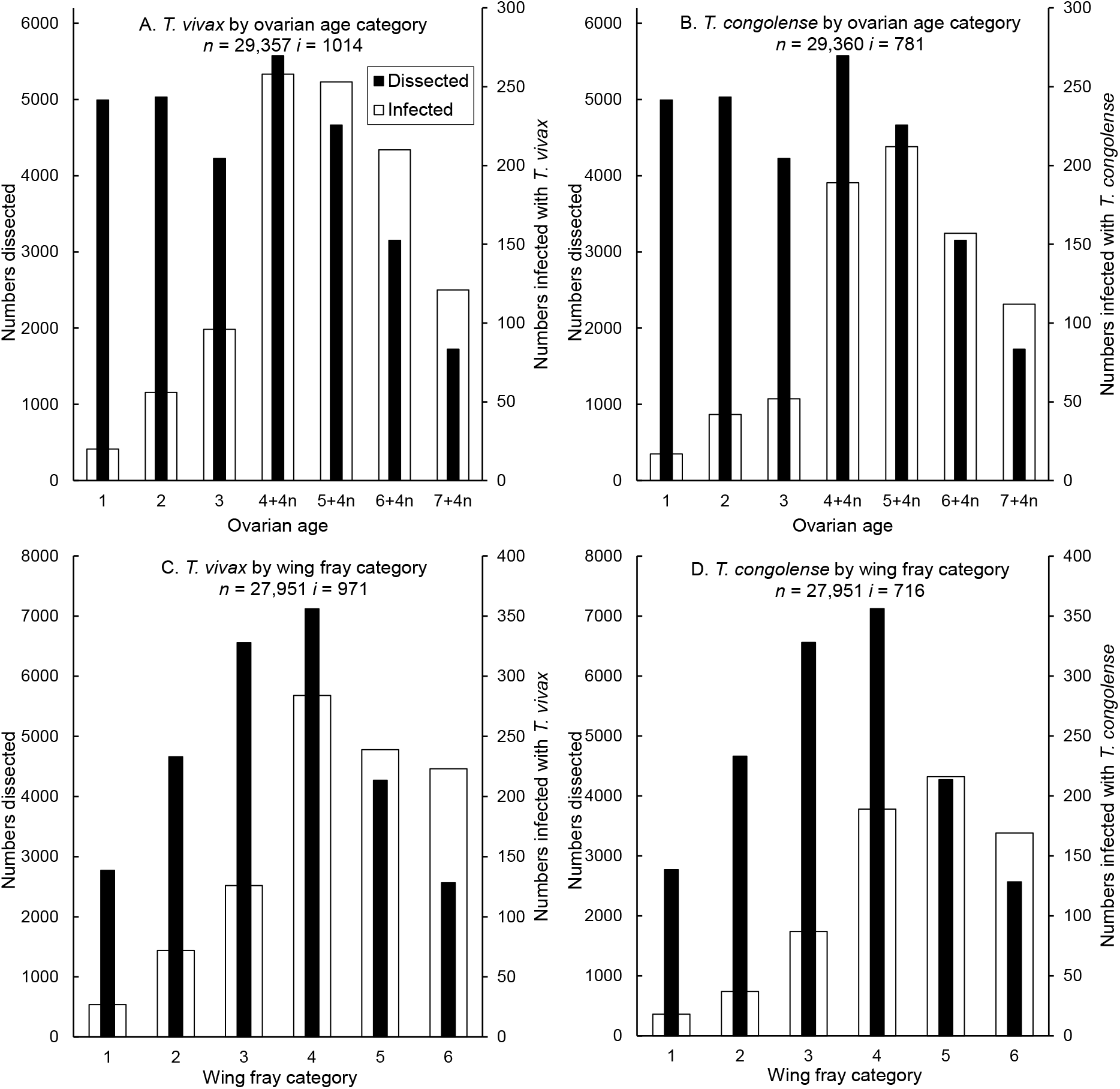
Distribution by age measures of female *G. pallidipes* captured at Rekomitjie Research Station 1990-1999. Figures show numbers (*n*) dissected and numbers (*i*) infected. **A., C.** *T. vivax*; **B., D.** *T. congolense* by ovarian age and wing fray categories, respectively. Notice the difference in scales on the left and right vertical axes on all graphs.

### Trypanosome prevalence as a function of fly age and ambient mean temperature

Stepwise logistic regressions were carried out, using the prevalences of either *T. vivax* or *T. congolense* infections as the dependent variable. Ovarian age, and wing fray, categories and mean ambient temperatures all accounted for significant fractions of the variance for both dependent variables. For both species of trypanosome, ovarian category removed the largest proportion of the variance, and the odds ratios increased steadily with age (Table 1). Once this variation was removed, the odds ratio for increases with increasing wing fray category were more modest.

**Table 1.** Results of logistic regression of the prevalence of *T. vivax* (A) and *T. congolense* (B) in female *G. pallidipes*, as a function of the fly’s ovarian, and wing fray, category and the mean daily temperature over the nine days prior to its capture. Sample size 27,951.

|  | Odds ratio | 95% Conf. Interval |
| --- | --- | --- |
| tbar91 | 0.956*** | 0.937 - 0.976 |
| Ovarian category |  |  |
| 1 | 1.00 |  |
| 2 | 2.78*** | 1.63 - 4.73 |
| 3 | 5.43*** | 3.26 - 9.04 |
| 4 | 9.67*** | 5.90 - 15.87 |
| 5 | 10.83*** | 6.56 - 17.86 |
| 6 | 12.51*** | 7.53 - 20.77 |
| 7 | 12.64*** | 7.48 - 21.37 |
| Wing fray category |  |  |
| 1 | 1.00 |  |
| 2 | 1.17 | 0.75 - 1.84 |
| 3 | 1.01 | 0.66 - 1.56 |
| 4 | 1.32 | 0.87 - 2.02 |
| 5 | 1.60* | 1.04 - 2.46 |
| 6 | 2.42*** | 1.57 - 3.75 |
| Constant | 0.011 | 0.005 - 0.022 |

B. *T. congolense*
|  | Odds ratio | 95% Conf. Interval |
| --- | --- | --- |
| tbar91 | 0.949*** | 0.927 - 0.972 |
| Ovarian category |  |  |
| 1 | 1.00 |  |
| 2 | 2.69** | 1.45 - 4.99 |
| 3 | 3.49*** | 1.88 - 6.46 |
| 4 | 7.58*** | 4.23 - 13.58 |
| 5 | 9.68*** | 5.39 - 17.40 |
| 6 | 9.88*** | 5.44 - 17.92 |
| 7 | 11.76*** | 6.39 - 21.63 |
| Wing fray category |  |  |
| 1 | 1.00 |  |
| 2 | 0.93 | 0.52 - 1.64 |
| 3 | 1.11 | 0.66 - 1.89 |
| 4 | 1.37 | 0.81 - 2.31 |
| 5 | 2.22** | 1.32 - 3.76 |
| 6 | 2.76*** | 1.62 - 4.71 |
| Constant | 0.010 | 0.004 - 0.023 |

Trypanosome prevalences, adjusted for ovarian and wing fray categories, declined significantly with increasing temperature (Table 1) measured as the mean temperature averaged over the nine days prior to the day on which the fly was captured – this period approximating the duration of the pregnancy cycle. The result is in accord with the finding that trypanosome prevalences, pooled on age, generally decrease in the hottest months of the year (Figure 1B).

### Fitting data using the simple catalytic function

The model defined in Equation (1), based on the assumption that no flies ovulate more than seven times, provides a reasonable fit to the data for both species of trypanosome (Figure 3) – although there are clear trends in the residuals. For *T. vivax*, the best fit was obtained with λ = 0.00141 per day and τ = 9.7 days (Table 2), very close to the values of λ = 0.00149 per day and τ = 14.2 days of Woolhouse *et al*. (1993), using only data for flies dissected in 1990 and 1991. For *T. congolense*, using the same model, the values were λ = 0.00107 per day and τ = 9.8 days (Table 2). There is considerable overlap of the 95% confidence intervals between species, for λ and for τ. Results obtained using our larger data set also suggest that there is no need to postulate a decline in *T. congolense* prevalence among older flies (Figure 3B, *cf*. Woolhouse *et al*., 1993).

**Figure 3.**
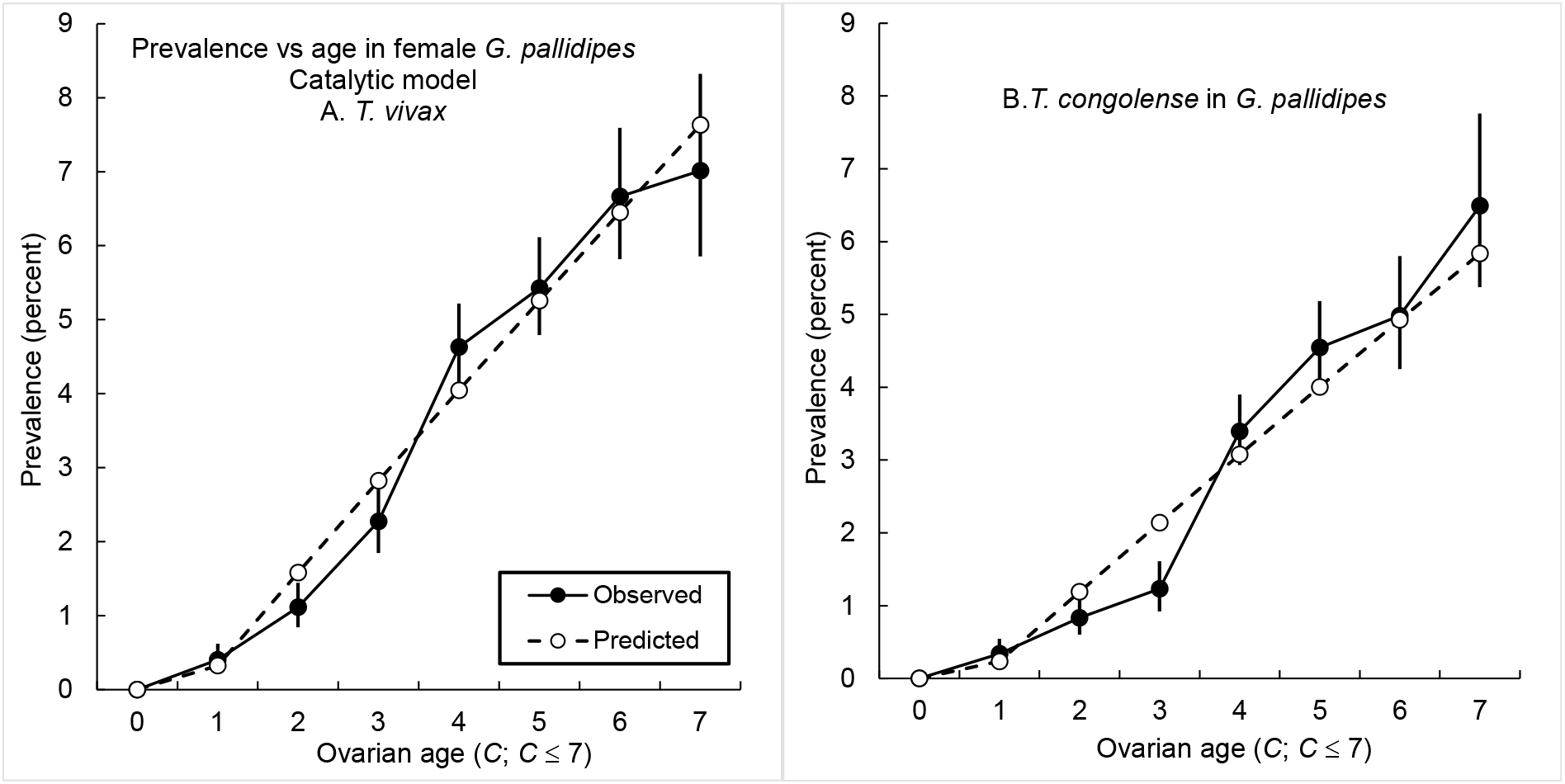
Fitting the function *P* (*a*) = 1 - exp [-λ(*a* - τ)] to age-specific prevalences for **A.** *T. vivax* and **B.** *T. congolense* in female *G. pallidipes* – where *a* is the age of the fly, λ the force of infection and τ the delay between the time that a fly is infected and when trypanosomes can first be detected in the fly. It is assumed that negligible numbers of flies survive >7 ovulations. Error bars on observed prevalences are exact 95% confidence interval for a binomial proportion.

**Table 2.**
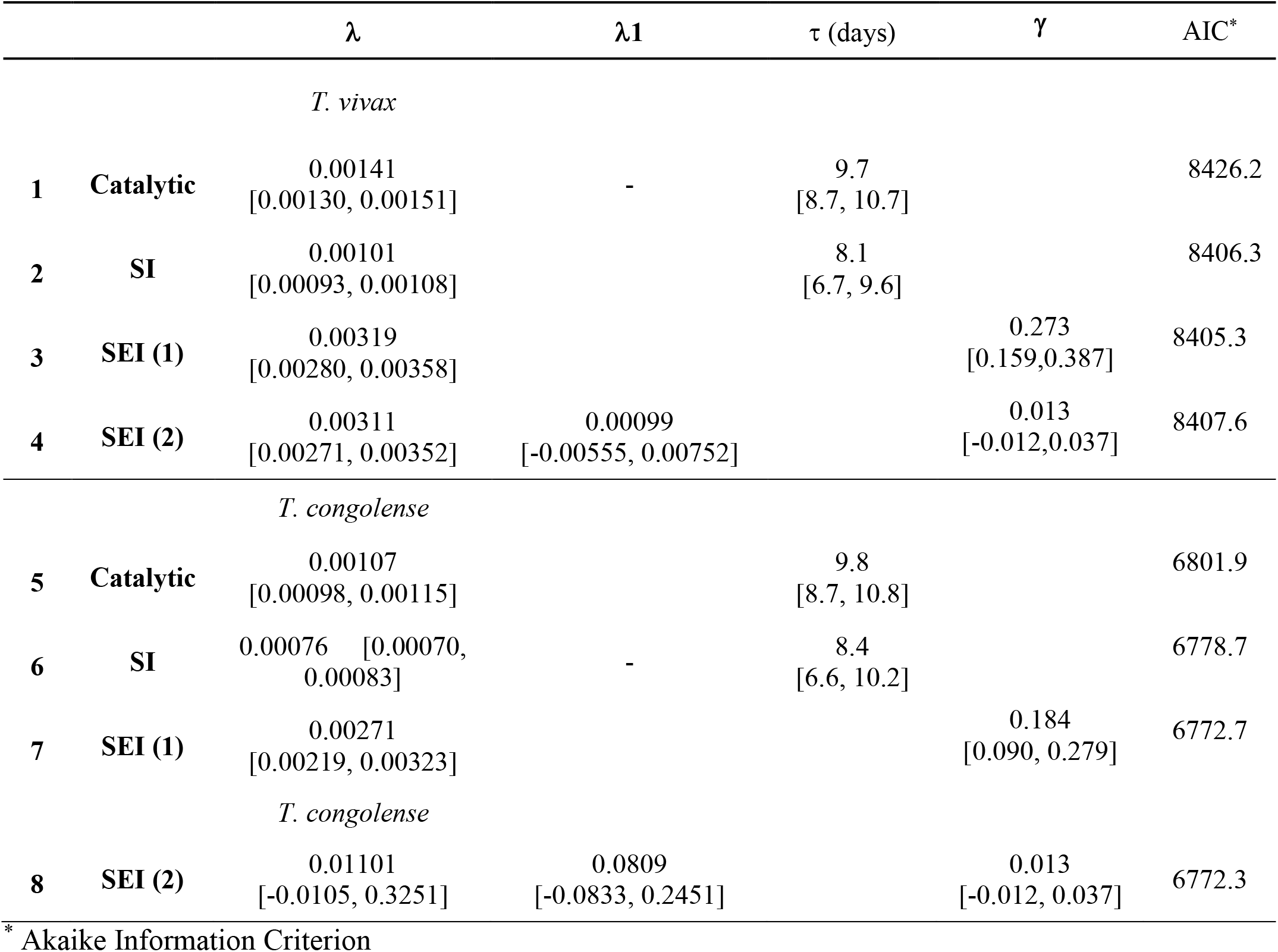
Maximum likelihood estimates for catalytic, SI, and SEI infection models, for *T. vivax* and *T. congolense* infections of female *G. pallidipes*. For the SEI(2) model the force of infection (λ1) at the first meal can be different from that at all subsequent meals. For all other models λ is identical for all meals. Units are daily for the catalytic and SI models, and 3-day for SEI. Figures in parentheses below the estimate provide the 95% confidence interval (CI). We assume a fixed survival probability of 0. 930 /3da = 0.976/da, previously estimated via ML, from total count data only. If a CI spans 0, then that parameter is not significantly different from 0, when that parameter is added to a model already containing the other parameters. Lower AIC values within each species denote higher quality of model fit.

| | | $\lambda$ | $\lambda_1$ | $\tau$ (days) | $\gamma$ | AIC* |
| --- | --- | --- | --- | --- | --- | --- |
| *T. vivax* |  |  |  |  |  |  |
| 1 | Catalytic | 0.00141<br>[0.00130, 0.00151] | - | 9.7<br>[8.7, 10.7] |  | 8426.2 |
| 2 | SI | 0.00101<br>[0.00093, 0.00108] |  | 8.1<br>[6.7, 9.6] |  | 8406.3 |
| 3 | SEI (1) | 0.00319<br>[0.00280, 0.00358] |  |  | 0.273<br>[0.159,0.387] | 8405.3 |
| 4 | SEI (2) | 0.00311<br>[0.00271, 0.00352] | 0.00099<br>[-0.00555, 0.00752] |  | 0.013<br>[-0.012,0.037] | 8407.6 |
| *T. congolense* |  |  |  |  |  |  |
| 5 | Catalytic | 0.00107<br>[0.00098, 0.00115] |  | 9.8<br>[8.7, 10.8] |  | 6801.9 |
| 6 | SI | 0.00076 [0.00070, 0.00083] | - | 8.4<br>[6.6, 10.2] |  | 6778.7 |
| 7 | SEI (1) | 0.00271<br>[0.00219, 0.00323] |  |  | 0.184<br>[0.090, 0.279] | 6772.7 |
| *T. congolense* |  |  |  |  |  |  |
| 8 | SEI (2) | 0.01101<br>[-0.0105, 0.3251] | 0.0809<br>[-0.0833, 0.2451] |  | 0.013<br>[-0.012, 0.037] | 6772.3 |

\* Akaike Information Criterion

Examination of the raw data on which the prevalences were calculated shows, however, that the Woolhouse *et al*. (1993) assumption that a negligible fraction of flies survives to a third ovarian cycle cannot be sustained (Figure 2). The increase, between ovarian categories 3 and 4+4*n*, in the total numbers of flies caught and dissected, and the large jump in the numbers of infected flies – make it obvious that significant proportions of flies do survive to a third (or even later) ovarian cycle: and we need to account for this fact. Note that a jump in observed trypanosome prevalence is not predicted by the fitted catalytic model (Figure 3).

### Fitting data using a Susceptible-Infected (SI) model

The SI model defined by Equations (3) and (4) requires knowledge of the survival probability. From the ovarian age distribution of the total number of adult female *G. pallidipes*, dissected to detect infections of *T. vivax* (Figure 2A), we estimated that Φ = 0.804, φ = Φ ^(1/9)^ = 0.976 and the daily mortality rate μ = -ln(φ) = 2.500%. Using this estimate, and the procedures leading to Equations (3) and (4), resulted in improved fits to the data for *T. vivax*, relative to the catalytic model (*cf* Figures 3A and 4A; AIC values (Table 2) differ by about 20).

**Figure 4.**
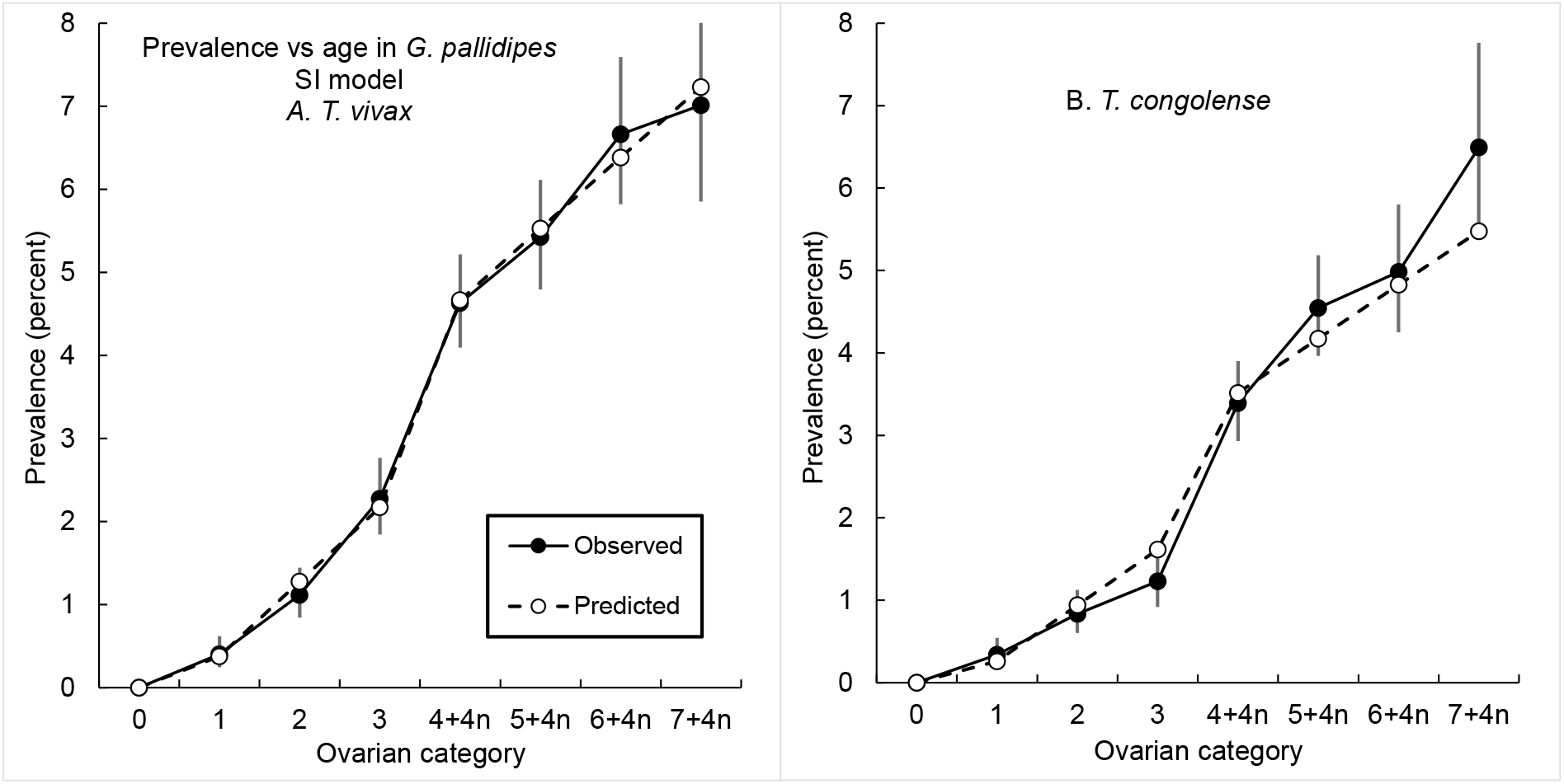
**A.** Fitting age-specific percentage prevalences of **A.** *T. vivax* and **B.** *T. congolense* in female *G. pallidipes*, using a Susceptible-Infected (SI) model. Prevalences were calculated from the average of the predicted numbers of infected and uninfected flies over successive 9-day pregnancy periods. It is assumed that negligible numbers of flies survive >15 ovulations. Error bars on observed prevalences are exact 95% confidence interval for a binomial proportion.

For *T. congolense*, similarly, the fit was markedly improved (*cf* Figures 3B and 4B), with the AIC values again differing by about 20. As is obvious from the inspection of Figure 4, however, the fit to the *T. vivax* data is markedly better than for *T. congolense* when the SI model is applied to the data for both species.

### Fitting data using a Susceptible-Exposed-Infected (SEI) model

The relatively poor fit of the SI model to *T. congolense* data results from the use of the value (τ) to allow for the time taken to mature an infection. Using the SEI model, described by Equation (5), results in an improved fit to the data for *T. congolense (cf* Figures 5A, 4B). The AIC for the SEI model was 6.0 lower than that for the SI model, Table 2, rows 4 and 5). For *T. vivax* the AIC criteria achieved using the SI and SEI models differed by only 0.4 < 2.0 so that there was no “quality of fit” difference between these two model forms (Table 2, rows 2 and 3). Visually, the fit obtained using the SEI model (not shown) is almost indistinguishable from that for the SI model (Figure 4A).

**Figure 5.**
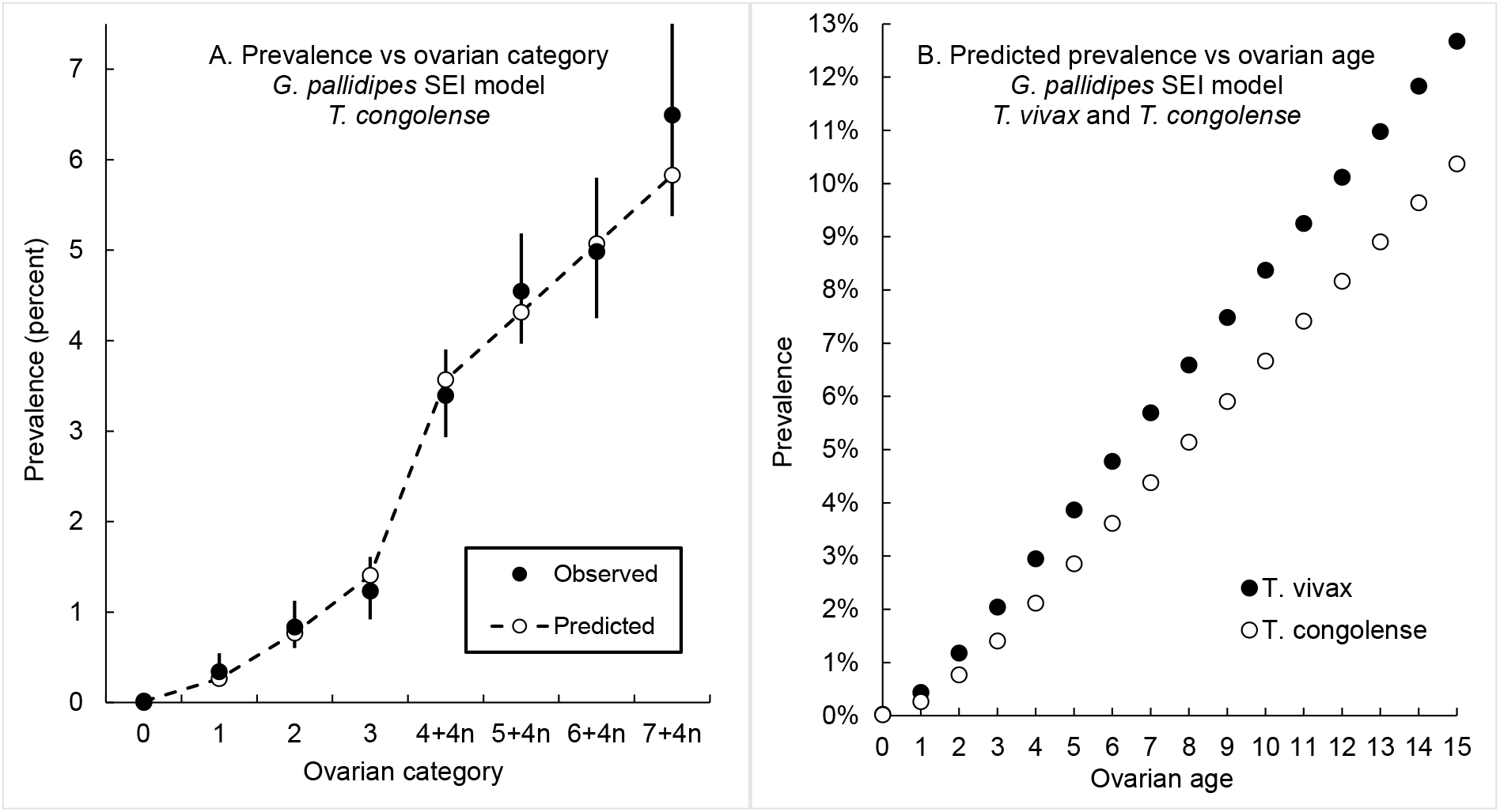
**A.** Fitting age-specific percentage prevalences of *T. congolense* in female *G. pallidipes*, using a Susceptible-Exposed-Infected (SEI) model. **B.** Predicted prevalence for ovulation ages 1-15, calculated from the average of the predicted numbers of infected and uninfected flies over successive 9-day pregnancy periods. It is assumed that negligible numbers of flies survive >15 ovulations. Error bars on observed prevalences are exact 95% confidence interval for a binomial proportion.

When we allowed the possibility that the force of infection for *T. congolense* was higher for the first bloodmeal (λ1) than for all subsequent meals (λ), the 95% CIs for the estimates of λ1, λ and γ all span 0, indicating that the teneral effect is not statistically significant in our data (Table 2, row 8). In further support of this, the AIC differed by <2.0, for both species of trypanosome, from the model form where we assumed λ1 = λ (Table 2, rows 3 vs 4 and 7 vs 8). Note also that, in single-lambda model, there was considerable overlap in CIs, between *T. vivax* and *T. congolense*, for the estimates of λ and γ (Table 2, rows 3 and 7). Thus, while the point estimates of the two parameters indicate a higher probability of infection for *T. vivax* – and a shorter maturation time, resulting in higher predicted prevalences for this species (Figure 4B), the difference is not statistically significant.

## Discussion

Recognition of the fact that significant proportions of female tsetse survive long enough in the field that they ovulate more than seven times means that it is not appropriate to use a simple catalytic model to fit age-prevalence data for trypanosome infections. The more complex SEI model provides much improved fits to the data but requires knowledge of the mortality rate among adult female tsetse. This can be a problem because estimating this mortality from age distributions has proved to be extremely difficult (Hargrove & Ackley, 2015;), particularly using samples collected over a short period of time. More believable estimates of mortality can be obtained, however, if samples are pooled over time periods of a year or more – such that problems associated with the seasonal instability of the population age structure are avoided (Van Sickle & Phelps, 1988; Ackley & Hargrove, 2017). Accordingly, we have only applied our analysis to data pooled over the whole period of the study and have not attempted to estimate the parameters λ, λ1 and γ over shorter periods of time.

### Coinfection rates

Coinfections, with more than one trypanosome species in a single fly, can be detected using techniques such as the polymerase chain reactions (PCR) that detect and identify, to species level, the DNA of different trypanosomes. The dissection process we used does not allow of such exact identification. Notice, however, that any fly that only has a mouthpart infection, can have only a *T. vivax*-type infection – because, if it also had a *T. congolense* infection, it would have trypanosomes in the midgut. The possibility of a *vivax*-*congolense* coinfection arises, therefore, only in flies where there are trypanosomes detected both in the mouthparts and the midgut. The following calculation suggests that the chances of this coinfection occurring for the female *G. pallidipes* in our study is negligible. Thus only 2.66% and 3.45% of all flies dissected were diagnosed to have a *T. congolense-type* or *T. vivax*-type infection, respectively. If a fly’s chances of acquiring an infection of one of the trypanosome species is independent of whether it is already infected with the other species, then the probability of finding a fly that is positive for both species is thus 0.0266 × 0.0345 = 0.00092 or about 1 in 1000 flies. Coinfections would be even rarer if, as seems likely, infection with one species of trypanosome reduced the probability of acquiring an infection with another trypanosome species. Coinfections could be more prevalent than indicated above only if an infection with one species of trypanosome *increased* the probability of acquiring an infection with the other species. We are not aware of any suggestion in the literature that his might happen.

### T. congolense infections: the teneral effect

Welburn & Maudlin (1992) found that laboratory *G. m. morsitans* were about 7-times more likely to acquire a *T. congolense* infection at their first meal than at any subsequent meal. By contrast, in *G. pallidipes* sampled in the field, we could not reject the null hypothesis of equality between the force of infection for *T. congolense* at the first vs all later meals. Given the very large sample size available to us, and the excellent fit of the SEI model to the data, we suggest that it might not even be feasible to separate λ1 and λ using field data.

The following calculations suggest that, in any case, a teneral effect of the order estimated by Welburn & Maudlin (1992) would be of little importance even if it did occur – basically because of the longevity of tsetse, particularly females, and their need to feed every 2.5-3 days (Randolph et al., 1992, Hargrove, 1999). Thus, suppose first that there were no teneral effect and that λ = 0.003 at every meal, including the first (Table 2, row 7). Then, if a fly lives for 120 days (Hargrove et al., 2010) we expect it to take about 40 blood meals. Its chances of remaining *uninfected* with *T. congolense*, throughout its life, is then estimated as (1 - 0.003)^40^ = 0.887, and the probability that it is infected is 1 – 0.887 = 11.3%, in good accord with the observed data and the predictions of our model (Fig. 5B). Suppose, alternatively, that there is a teneral effect of the order suggested by Welburn & Maudlin (1992), such that λ1 = 7λ = 0.021. Then the estimated probability of remaining uninfected for 120 days is (1-0.021) × (1 - 0.003)^39^ = 0.871 and the probability that the fly has become infected is 1 – .871 = 12.9%, only slightly higher than when there is no teneral effect. We thus find no support, at least in *G. pallidipes*, for Welburn & Maudlin’s (1992) suggestion that bloodmeals other than the first are of no importance epidemiologically and conclude, conversely, that the so-called teneral effect is of minor importance in the epidemiology of *T. congolense*.

### Trypanosome prevalence in vertebrate hosts

The probability (λ) that tsetse will mature a trypanosome infection after taking a bloodmeal is the product λ = *p*1 × *p*2, where *p*1 is the probability that the bloodmeal they take is indeed infected with trypanosomes, and *p*2 is the conditional probability that a fly matures an infection given that it takes an infected meal. Results for the SEI model (Table 2, row 7) suggest λ ≈ 0.003 per 3-day period (*i.e*., per bloodmeal) for *T. congolense* infections of female *G. pallidipes*. From Welburn & Maudlin (1992, Figure 1) *p*1 for the first bloodmeal is about 0.7 and *p*2 for all later meals is about 1/7 of this value, *i.e., p*2 = 0.1. If *p*2 takes a similarly low value for non-teneral female *G. pallidipes* at Rekomitjie, this suggests that the proportion *p*1 that take a meal from a host infected with *T. congolense* is *p*1 = λ/ *p*2 = 0.003/0.1 = 0.03. That is to say, of the hosts fed on by *G. pallidipes* at Rekomitjie, only about 3% are harbouring sufficient *T. congolense* to ensure that tsetse feeding off them take an infected meal. If that is the case then the proportion of flies developing a mature infection after their first meal is λ = 0.03 × 0.7 = 0.021, or about 2.1% of the population

For *T. vivax* our estimate for λ is also about 0.003 (Table 2, line 3) but we do not have any estimate for *p*2. If it is markedly higher than for *T. congolense* then the proportion of animals with *T. vivax* infections is even lower than for *T. congolense*. For example, if *p*2 = 0.5 then *p*1 = λ2/ *p*2 = 0.003/0.5 = 0.006 and we predict that only 0.6% of vertebrate hosts would be harbouring sufficient *T. vivax* to provide an infective meal. Such low levels of infection in wild hosts support our suggestion that the teneral effect is unlikely to be important in the epidemiology of either *T. vivax* of *T. congolense*; the chances of becoming infected by a single meal is simply too small to explain the observed age-related changes in trypanosome prevalence.

### Sex and species differences

In the current study we have considered only females of *G. pallidipes*, because there is no method available for gauging the chronological age of males. Nonetheless, the degree of wing fray in males does provide estimates of relative age, albeit with less accuracy than is associated with the use of ovarian dissection in females. We also have available less extensive data for age-specific trypanosome prevalences in female *G. m. morsitans* sampled at Rekomitjie. Future work will investigate whether similar models to those developed here for female *G. pallidipes* are also appropriate for the pattern of acquisition of trypanosome infection for male *G. pallidipes*, and for *G. m. morsitans*. Given that the teneral effect was suggested based on trypanosome infections acquired by *G. m. morsitans* in the laboratory, we need to decide, particularly, whether the effect is important in field populations of this tsetse species.

## Acknowledgements

The authors acknowledge the support and continued interest of Dr William Shereni, Director of the Trypanosomiasis Control Department, Government of Zimbabwe. All experimental work was carried out at Rekomitjie Research Station in the Zambezi Valley. Our grateful thanks are due to the tsetse capture and dissection teams, ably led by Mr. Pio Chimanga. We acknowledge the contribution of Prof Mark Woolhouse, who trained the original field team to dissect flies to detect trypanosome infections.

## Author contributions

JWH designed the study, collated the data and carried out preliminary analyses. JVS carried out detailed statistical analysis. Both authors contributed equally to the preparation of the manuscript and the preparation of all figures and tables.

## Ethical standards

Not applicable; the study does not involve mammalian subjects.

## Consent for publication

Both authors gave consent for this publication. No other consent required.

## Availability of data and materials

All data used, and materials generated, in this publication are available from the authors on reasonable request.

